# Nonnative Fish in the Pearl River Delta: A Comprehensive Study on Their Salinity Adaptability and Potential Distribution Area in Brackish Water

**DOI:** 10.1101/2024.01.16.575818

**Authors:** Du Luo, Dingtian Yang, Yuefei Li, Jie Li

## Abstract

After extensive aquaculture, urbanization, and industrial development, the Pearl River Delta (PRD) has undergone profound environmental changes, with biological invasion emerging as a consequential anthropogenic byproduct, carrying significant implications for natural ecosystems, human health, and the economy. Despite these transformations, our understanding of nonnative fish species and their adaptability to the dynamic environmental conditions within the intricate river network of the PRD remains incomplete. To address this, we conducted a systematic survey of nonnative fish species in PRD waters through literature reviews. Then we analyzed the salinity adaptation of these fish, coupled with a spatial examination of salinity distribution, to comprehensively understand the potential invasion mechanisms of exotic fish in the region. Our findings identified 25 nonnative fish species in PRD waters, with three being translocated species within China and the remaining 22 being exotic introductions. Phylogenetically, these fish are broadly distributed among several orders, with Cichliformes, Perciformes, Cypriniformes, and Siluriformes emerging as the predominant orders. Despite most of the nonnative fish being freshwater species and exhibiting species-specific differences in salinity adaptability, they generally demonstrated an average salinity tolerance of 23.74 ppt, allowing them to adjust to brackish water. The diversity of species declined as the relative distance decreased from the upstream of the Pearl River Estuary (PRE) to its mouth in the sea (p < 0.01). Spatial analysis unveiled pronounced heterogeneity in salinity distribution, particularly in the east PRE of the LingDing Bay, featuring a sizable area with brackish water. Notably, under salinity intrusion circumstances, a gap region between salinity levels of 5.0 ppt and 15.0 ppt, spanning 3111.2 km2, emerged as a potential adaptation zone for nonnative fish. By emphasizing salinity as a key factor, our study may contribute to elucidating invasion mechanisms and enhancing the assessment of invasion risks in estuarine environments.

## 1. Introduction

Biological invasion represents one of the five major environmental challenges facing the world today and are integral to global change [1]. Consequently, invasive species are considered among the most serious ecological and economic threats [2]. The issues they cause, including the reduction of biodiversity, impacts on economic development, and threats to public health, have garnered widespread international attention. Estuarine areas are among the most heavily impacted zones by human activities, leading to significant ecological changes and a sharp decline in biodiversity. In particular, invasive species contribute significantly to the loss of diversity in estuarine environments [3]. The preservation of ecosystem functions and biodiversity in estuarine regions is increasingly recognized as crucial, highlighting the importance and urgency of research on invasive species in estuarine waters.

The estuarine ecosystem is one of the ecosystems at the highest risk of invasion worldwide, with numerous invasive species having become main members of estuarine ecosystems [4]. Fish are the most commonly introduced aquatic group, and the invasion of nonnative fish is considered a major contributor to the decline in native aquatic biodiversity [5]. Meanwhile, salinity and dissolved oxygen are believed to be the primary shaping factors for the specific ecological characteristics of estuarine environments, and differences in environmental factors may lead to fish adopting corresponding competition and predation strategies [6]. Invasive fish with strong salinity adaptability possess the ability to invade both freshwater and coastal marine environments [7]. Studies indicate that non-indigenous freshwater fish are more likely to invade estuarine areas, highlighting the critical role of salinity adaptability in enhancing their invasion capability in brackish water environments [8].

In a global context, the abundance of fish species within estuaries is shaped by spatially hierarchical predictors, with their richness influenced by the broader distributions and habitat utilization patterns of marine and freshwater species capable of colonizing estuaries [9]. Estuarine ecosystems constitute essential habitats for the completion of life cycles by estuarine fish species. Changes in species richness may be correlated with extreme tidal fluctuations and elevated water salinity [10]. The habitat features, along with seasonal fish movements associated with salinity intrusion, could contribute to the development of more diverse fish assemblages within the transitional zone of an estuarine–freshwater ecosystem [11]. Estuarine fishes can be categorized into various functional types, such as marine, estuarine, diadromous, and freshwater types [12]. Despite variations in the ecological types of dominant fish species in estuarine ecosystems across different regions [13], the distinctions in species within estuarine waters may play a crucial role in influencing ecosystem service functions [14]. The salinity disparity across various water areas within estuaries can be considerable, and the interplay between salinity levels and topographic characteristics may exert influence on the species richness of invasive fish [15]. Previous research has identified a negative correlation between freshwater discharge and the size of fish species [16]. Substantial variations in both the quantity and richness of fish species have been noted among distinct river mouths within the same geographic region [17]. The distribution of estuarine fish exhibits both temporal and spatial variations, with salinity fluctuations identified as a primary factor influencing the restructuring of fish communities [18]. Saltwater intrusion and hydrological alterations have the potential to induce the invasion of exotic fish species into estuarine ecosystems [19,20]. Furthermore, marine fish may also encroach upon estuarine waters [21].

The Pearl River Estuary (PRE) stands out as a crucial region of land-sea interaction in the northern South China Sea, hosting a diverse range of tropical and subtropical organisms. This area plays a vital role in supporting the feeding, migrations, and spawning activities of numerous aquatic species [22]. Despite its ecological significance, recent studies have revealed a concerning decline in fish biodiversity within the Pearl River Delta (PRD). Comparing current findings to records from the 1980s, there has been a 45% reduction in the number of fish species [23]. Furthermore, certain areas are experiencing an uptick in the number of nonnative fish species and the prevalence of major invasive species [24]. The diversity of invasive fish species and the dominance of key invasive species have shown an upward trend. Adding to the complexity, the PRD is a hub for the import and export of ornamental fish species [25]. Despite the potential exacerbation of invasive fish in estuarine waters, there is currently a significant research gap concerning the invasion mechanisms of nonnative fish species in the PRE. This underscores the need for further investigation into this understudied aspect of the region’s aquatic ecology.

Examining the compilation of nonnative fish species and delving into their adaptability to the primary environmental factor of salinity in the PRD could shed light on the processes and mechanisms underlying fish invasion in estuarine waters [26]. In this study, we systematically investigated and compiled information on nonnative fish species existing in the natural waters of the PRD. Additionally, we integrated available research data to analyze their tolerance and adaptability to brackish waters. Employing simulated analyses of salinity distribution patterns during salinity intrusion in dry-season scenarios, we conducted a spatial analysis to evaluate the potential adaptive capacity of nonnative fish in the PRD.

## 2. Materials and Methods

### 2.1. Study area

The Pearl River extends 2214 km, and has a drainage area of 452, 000 km^2^ in Southern China. It flows into the northern South China Sea via eight outlets of Humen, Jiaomen, Hongqimen, and Hengmen; Modaomen, Jitimen, Hutiaomen, and Yamen in the PRD [27] (Figure 1). Modaomen is the largest discharge and Humen is the second largest river inlet [28]. The LingDing Bay, receiving 53% of the river discharge, is the largest sub-estuary of the Pearl River [29]. The Pearl River is one of the three major drainage systems supplying the northern South China Sea [30]. The PRD is one of the most complex river systems in the world and the changes in shoreline and bathymetry caused by anthropogenic activity in the past several decades have induced variations on salinity distribution, stratification, and salt transport [31]. As a funnel-shaped estuary, most of the water depth in the PRE is shallower than 5 m, except for the two main channels in LingDing Bay with depths of 5–20 m [32]. Affected by both river discharge and the tides, the hydrodynamics of currents in the PRD are complicated. The geographic range used to calculate areas of the PRD was defined according to the Integrated Planning of Ecological Security System in the PRD (https://www.gd.gov.cn/gkmlpt/content/0/143/ post_143525.html#7; Accessed on 12 November 2023).

**Figure 1.**
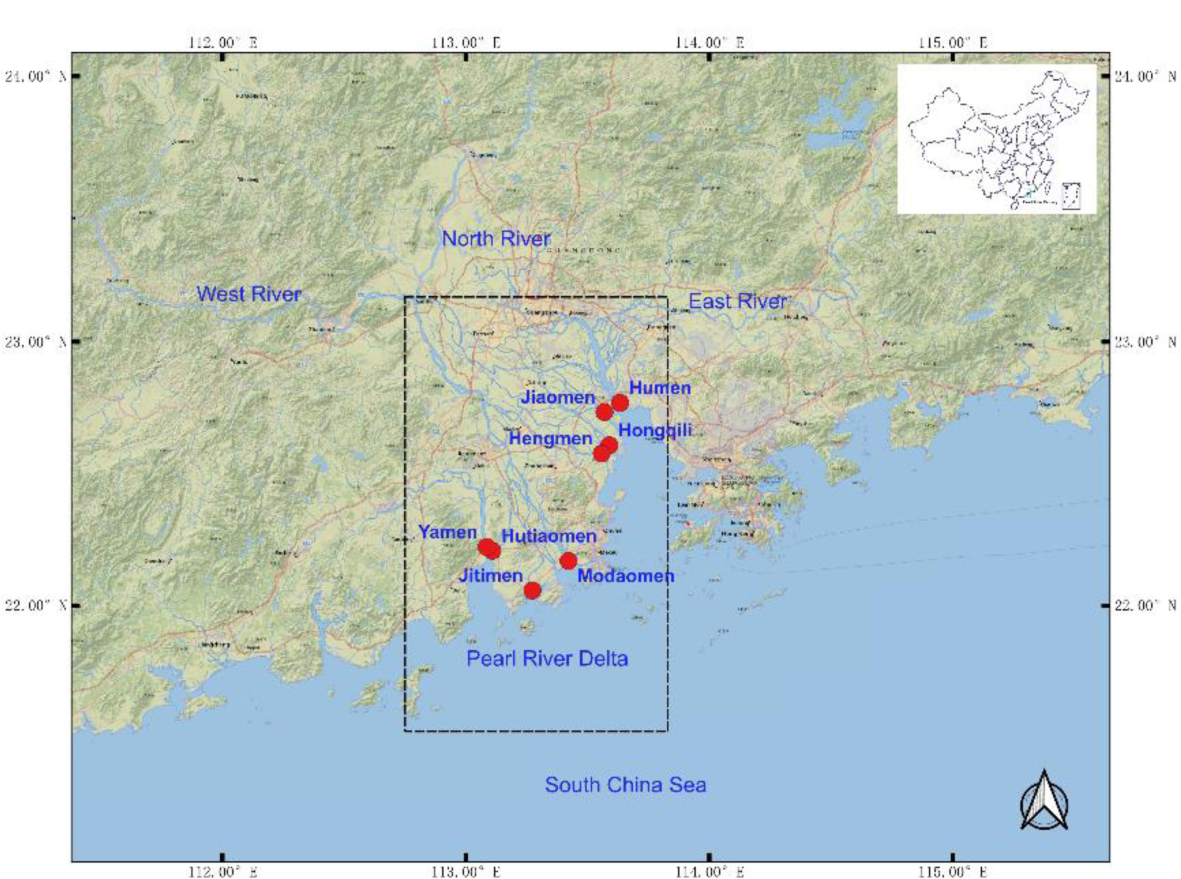
Geographic Location of the PRD.

### 2.2. Nonnative fish species and their salinity adaptation

Drawing from prior studies on nonnative fish invasion and fishery resource assessments in the Pearl River Basin [24,33–39], supplemented by our routine observations during fish resources investigation, we compiled a comprehensive dataset comprising sampling locations and the count of alien species. The introduction purposes for these nonnative fish were categorized into four groups: aquaculture, aquarium, both aquaculture and aquarium, and biocontrol. We assessed the taxonomic richness at the class, order, family, and genus levels for these exotic fish. To investigate the salinity adaptation of these nonnative fish, we conducted a thorough review of salinity tolerance for each species, with a particular emphasis on larvae. The semi-lethal concentration (LC50) within 96 hours or the maximum salinity recorded for fish survival, as documented in the references, was used to establish salinity limits. The salinity of typical seawater, recorded as 35.0 ppt, served as a reference point. The relative distance between each sampling area in the previous investigation and the pure seawater line in the epipelagic zone outside the mouth of the PRE was calculated using an online map calculator (https://map.baidu.com/; accessed on November 15, 2023). The trophic level for each fish species was determined based on information from FishBase (https://fishbase.se; accessed on November 15, 2023).

### 2.3. Water salinity data sources and spatial distribution

To simulate the scenario of salinity intrusion, we collected and screened available detected data with higher salinity during the dry season to construct a dataset. Salinity data for the Pearl River were obtained from publicly accessible sources, including the Department of Ecology and Environment of Guangdong Province (http://gdee.gd.gov.cn/; accessed on February 7, 2023) and the Environmental Protection Department of the Government of the Hong Kong Special Administrative Region (https://www.epd.gov.hk/; accessed on November 14, 2023). Additionally, we gathered information from published papers [40–42] and published books [36]. To mitigate potential biases induced by seawater in the interpolation process, water quality monitoring data from Jiangmen (http://www.jiangmen.gov.cn/bmpd/jmssthjj/index. html; Accessed on November 21, 2023) and Dongguan (https://dgepb.dg.gov.cn/; Accessed on November 21, 2023), situated on the outskirts of the PRD, were utilized as reference data. If the public water quality did not exceed preset standards, a salinity value of < 0.5 ppt was considered. The spatial analysis of salinity distribution was conducted using the open-source software QGIS version 3.34.

### 2.4. Statistical Analysis

The overall characteristics of salinity tolerance were examined using frequency distribution. Linear relationships between the relative distance to the shoreline of seawater and species richness, as well as the average trophic level, were analyzed through regression using GraphPad Prism version 9.0 with a 95% confidence level. The estimation of potential habitat area relied on simulated salinity level lines within the PRD range as defined previously. Spatial analysis interpolation was conducted using the Gaussian method in the Smart-Map plugin within the open-source software QGIS version 3.34 [43].

## 3. Results

### 3.1. Nonnative fish species in the PRD

The comprehensive survey revealed a total of 25 nonnative fish species inhabiting the natural waters of the PRD (Table 1). Among these species, Mrigal carp (*Cirrhinus mrigala*) and Nile tilapia (*Oreochromis niloticus*) have emerged as dominant species based on fishery resource investigations. Wuchang bream (*Megalobrama amblycephala*), Yangtze icefish (*Neosalanx taihuensis*), and striped snakehead (*Channa striata*) represent translocated species within China, while the remaining species are aliens introduced from foreign countries. The primary purposes for their introduction were aquaculture, followed by aquarium and biocontrol applications. Considering their utilization, over 20 species were cultivated as aquaculture fish, while 7 species were maintained as aquarium fish (Figure 2A). Three species, namely Mississippi paddlefish (*Polyodon spathula*), mirror carp (*Cyprinus carpio* var.), and striped snakehead (*Channa striata*), were employed both as food fish and as pet fish. The mosquitofish (*Gambusia affinis*) was historically introduced and utilized for mosquito control. Based on phylogenetic classification, these species belonged to 22 genera, 15 families, and 9 orders within the class Actinopterygii. In terms of order classification, Cichliformes (20%), Perciformes (20%), Cypriniformes (16%), and Siluriformes (16%) collectively constituted 72% of the nonnative fish species (refer to Figure 2B).

**Figure 2.**
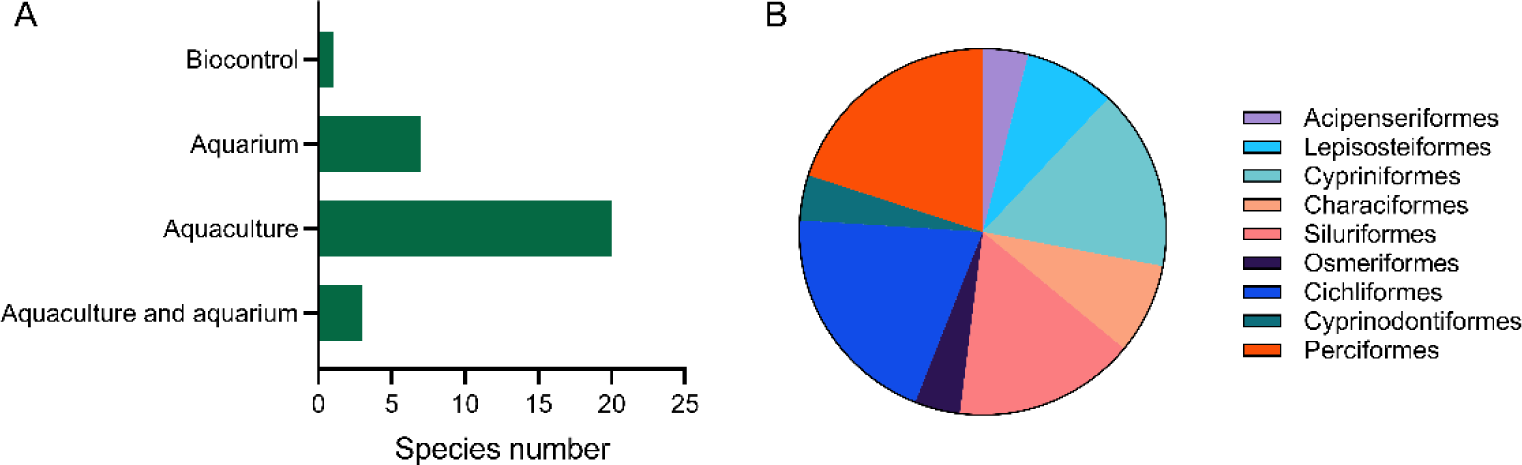
Introduction purposes for nonnative fish species in the PRD (A) and phylogenetic classification by order (B).

**Table 1.**
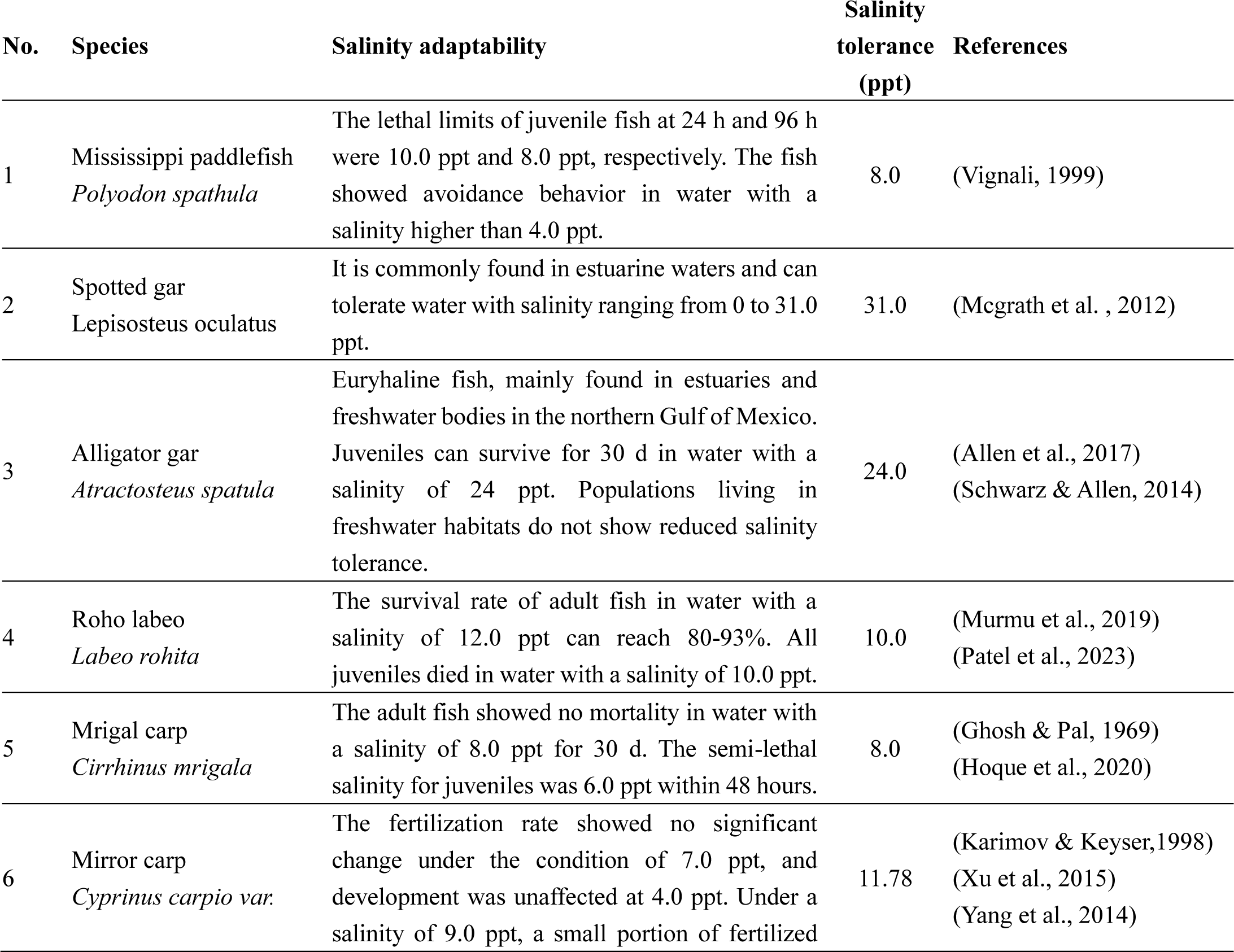
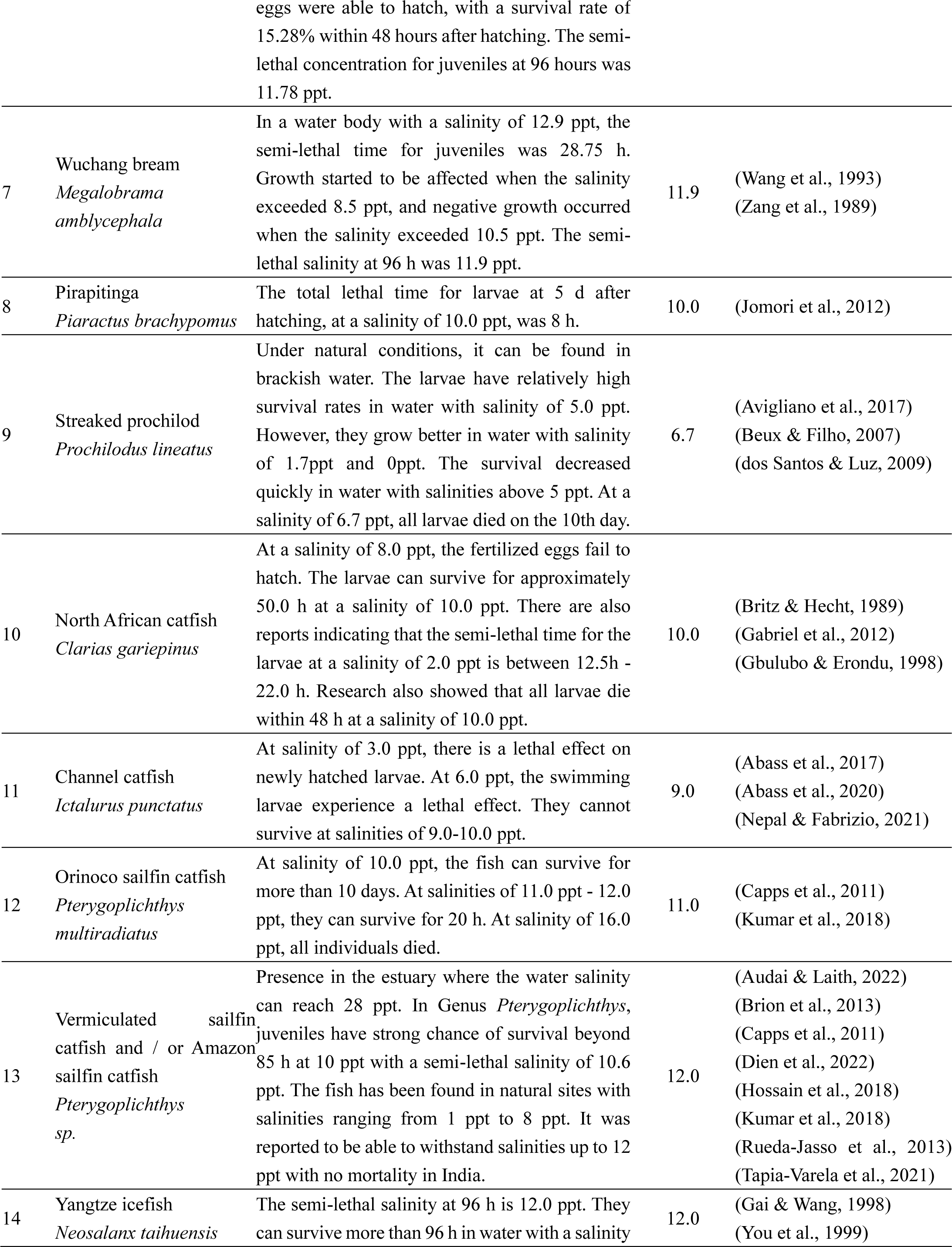
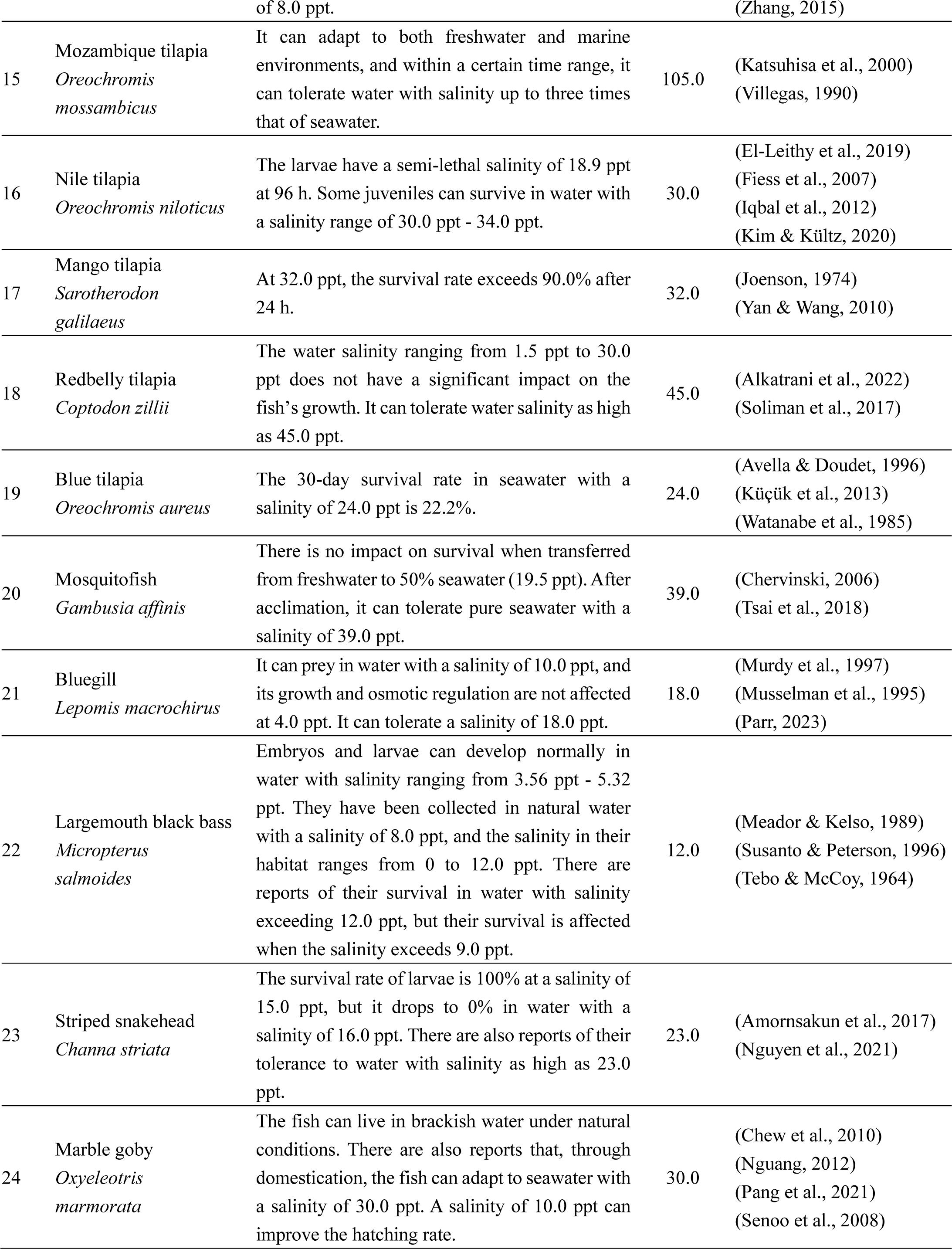
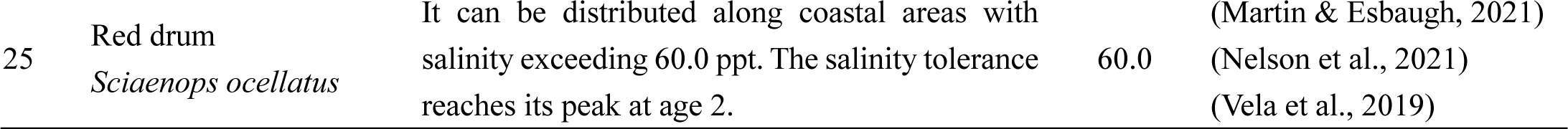
Nonnative fish species in the PRD and their salinity adaptability.

### 3.2. Salinity tolerance of the nonnative fish in the PRD

The results revealed a wide range of salinity tolerance among nonnative fish, with *Prochilodus lineatus* exhibiting the lowest tolerance at 6.7 ppt and *Oreochromis mossambicus* demonstrating the highest tolerance, three times that of marine water, averaging at 23.7 ppt (Figure 3A). Frequency distribution statistics indicated a median salinity of 12.0 ppt, with a 25th percentile at 10.0 ppt and a 75th percentile at 30.5 ppt (Figure 3B). Given that more than half of the fish’s salinity limits clustered around 10.0 ppt, the tolerance distribution of these nonnative fish species is non-discrete. Notably, species such as spotted gar (*Lepisosteus oculatus*), alligator gar (Atractosteus spatula), mosquitofish (*Gambusia affinis*), striped snakehead (*Channa striata*), and marble goby (*Oxyeleotris marmorata*) are typically considered freshwater fish. However, they exhibit strong salinity tolerance, ranging from 23.0 ppt to 39.0 ppt. Additionally, the five tilapia species originating from Africa demonstrated even higher salinity tolerance, with an average limit of 47.2 ppt. The red drum (*Sciaenops ocellatus*), classified as a marine fish, displayed a maximum salinity tolerance of 60.0 ppt.

**Figure 3.**
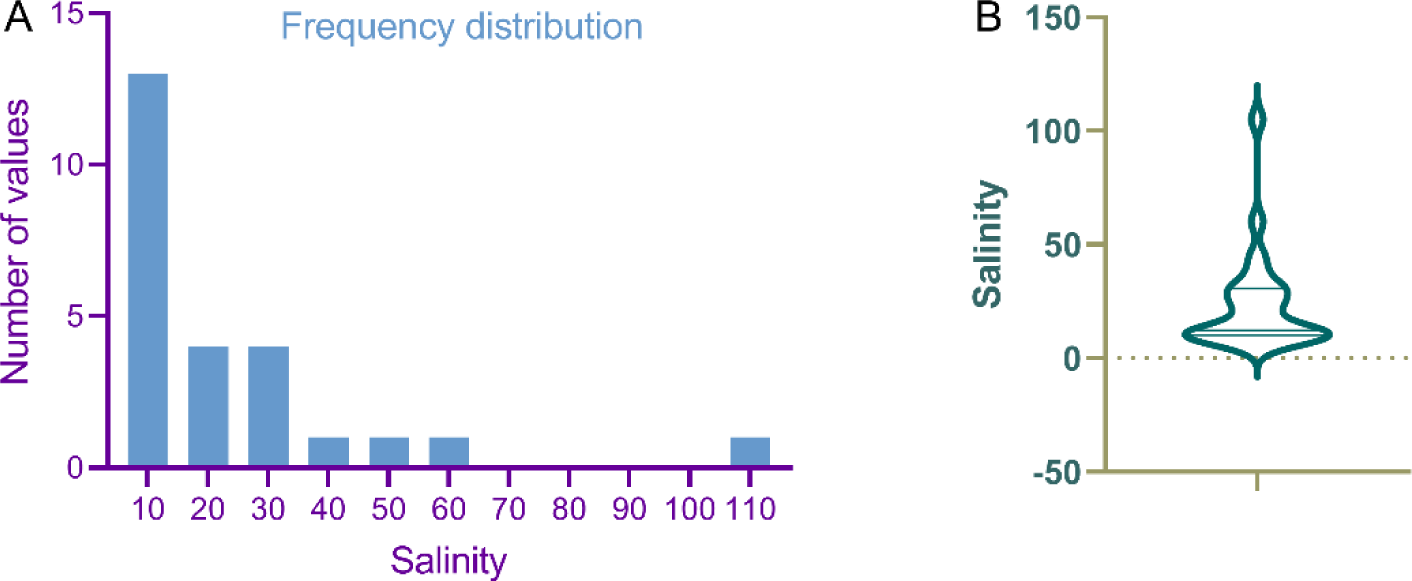
Frequency distribution of salinity tolerance in nonnative fish. The bar chart (A) depicts the number of salinity limit values distributed across saline gradients, while the violin chart (B) illustrates the aggregating characteristics.

### 3.3. Spatial variation in alpha diversity and trophic levels

The number of nonnative species exhibited variation across different geographic areas within the PRD. In Lingding Bay, located to the east of the PRE, common catches in fishery resources on both the east and west coasts include Mrigal carp (*Cirrhinus mrigala*), North African catfish (*Clarias gariepinus*), tilapia (*Oreochromis sp.*), and sailfin catfish. Linear analysis of geographic correlation revealed a significant decrease in the alpha diversity of nonnative fish species from the head of the PRD to the mouth of the PRE (R^2^ = 0.79, P < 0.0001) (Figure 4A). Data from one investigation indicated that the average trophic level index of nonnative fish in the PRD is 2.89 ± 0.79 se (n=15), ranging from the lowest at 2.0 ± 0.0 se in Nile tilapia (*Oreochromis niloticus*) to the highest at 4.2 ± 0.3 se in Channel catfish (*Ictalurus punctatus*). Consequently, linear relationship analysis revealed that the trophic level of alien species was higher in the lower reaches of the estuary, although no significant correlation was detected (R^2^ = 0.33, P = 0.06) (Figure 4B).

**Figure 4.**
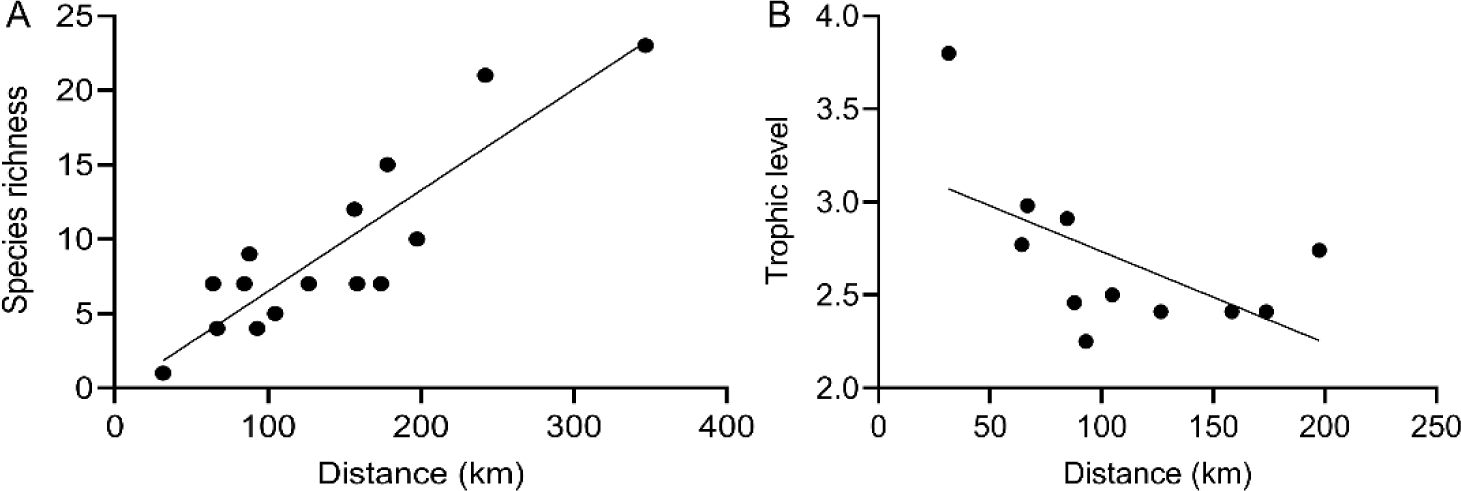
Simple linear regression demonstrating alpha diversity (A) and trophic level (B) variation relative to the distance of investigation areas from the seawater shoreline at the mouth of the PRE.

### 3.4. Spatial analysis of salinity distribution

In the estuarine zone where freshwater meets seawater at the PRE, there are highly pronounced spatiotemporal variations in salinity. Upstream in the estuary, the river network is intricate, and freshwater discharge changes with the seasons and tributaries. In the downstream areas of the estuary, the Pearl River connects to the South China Sea through three sub-estuaries: Lingding Bay, Modaomen, and Huangmaohai, forming an even more complex network of salinity variations in the delta [44]. Modaomen to the west experiences a significant influx of freshwater flow, causing drastic fluctuations in salinity at the convergence zone. On the eastern side, Humen features a longer brackish water mixing zone. Within the Lingding Bay, an area of approximately 2000 km² forms, extending about 70 km from the mouth to the Humen outlet, with an approximate width of 35 km at the LDB mouth [45]. Meanwhile, in the intermediate area of Hengmen, the flow of freshwater runoff runs almost perpendicular to the direction of the tidal flow, resulting in its salinity being the lowest among the eight estuary mouths.

A total of 132 sampling sites, selected from collections, were utilized in the simulation of salinity intrusion. Under the conditions of salinity intrusion, the front of the estimated 5.0 ppt line extended northward up to 22.85 N along the Humen inlet (Figure 5). Spatially, over 50% (9,827.3 km^2^) of the PRD experienced salinity levels exceeding 5.0 ppt. The area between 5.0 ppt and 10.0 ppt covered 2,021.8 km^2^, while the area between 10.0 ppt and 15.0 ppt encompassed 1,089.4 km^2^. The combined area between 5.0 ppt and 15.0 ppt accounted for 17.50% of the total PRD area. Brackish water with salinity < 15.0 ppt and freshwater collectively constituted 62.23% of the PRD area. Considering distances and areas, the majority of salinity intrusion occurred in the eastern part of the PRD, specifically in Lingding Bay along the Humen inlet.

**Figure 5.**
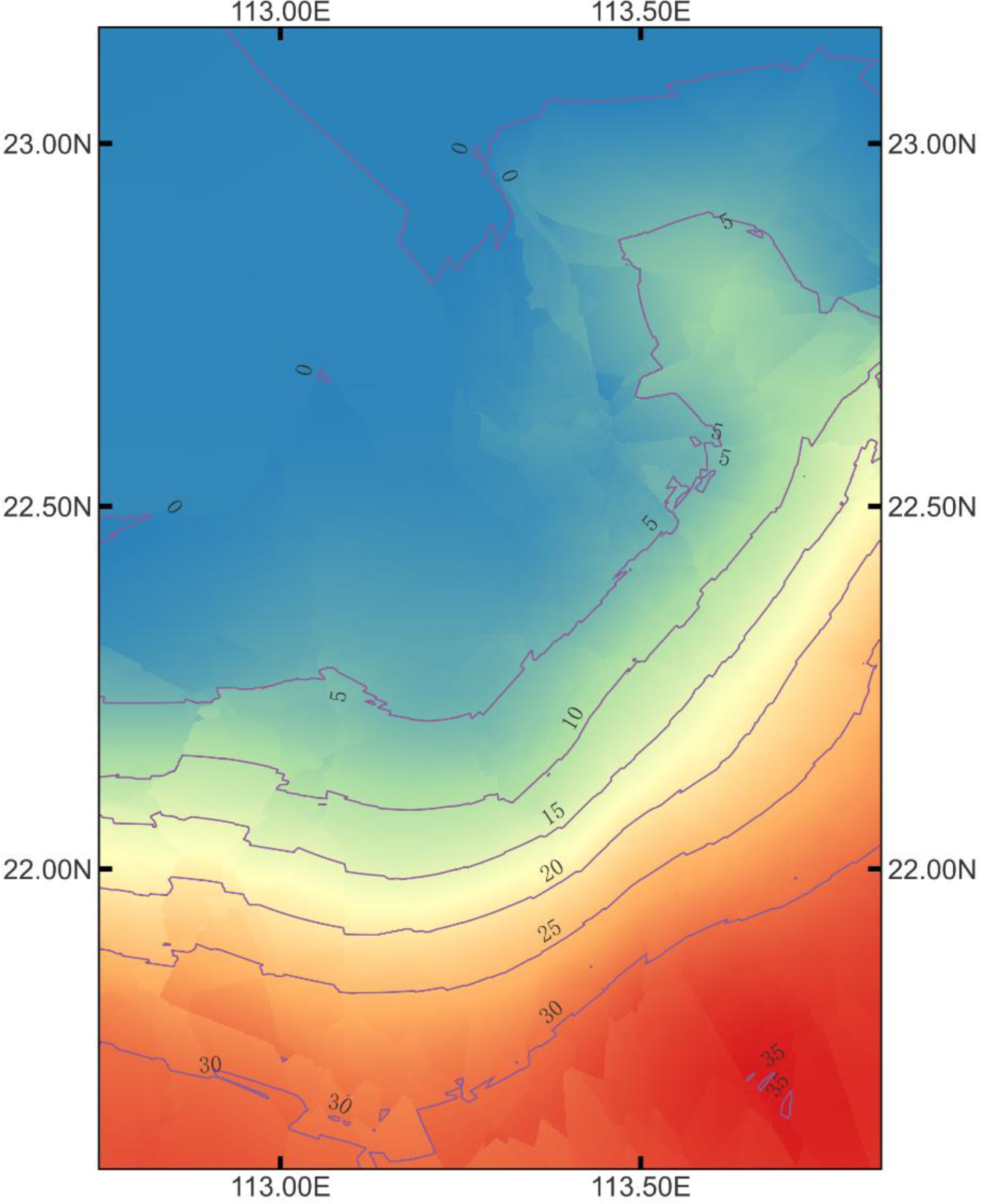
Simulated salinity distribution in the PRD. The map was created using the open-source software QGIS version 3.34. Interpolation was carried out using the Smart-Map plugin with the Gaussian method.

## 4. Discussion

The estuaries around the world are experiencing both biological and non-biological changes [46], highlighting the complexity of estuarine ecosystems. Climate change affects the timing of juvenile fish entering estuaries [47]. Additionally, climate warming weakens constraints on the spread of marine fish [48]. Changes in the population of bait organisms in local aquatic ecosystems undoubtedly influence the invasion of non-native fish species [49]. Against the backdrop of global changes, the environment in the PRD has undergone complex and substantial transformations. The PRD has experienced extensive anthropogenic activities during its remarkable economic development and urbanization [50]. A report in 2016 assumed that the habitat loss in PRE would reduce its carrying capacity. Exploratory simulations indicated that up to approximately 30.5% of the current habitat may be lost over the next two to three decades [51].

This study found that there are at least 25 species of exotic fish commonly found in the waters of the PRD, and some of these exotic fishes, such as Mrigal carp (*Cirrhinus mrigala*) and Nile tilapia (*Oreochromis niloticus*), have become widely distributed dominant species. In contrast, the fisheries resources of the PRD waters have shown a trend of decline. The fish assemblage structure has obviously changed in the past three decades [23]. Therefore, the changes in the structure and abundance of fish resources in the PRE are likely to be closely related to the invasion of exotic fishes. Similarly, in the San Francisco estuary, which is also affected by the invasion of exotic fishes, many species of fish have shown a downward trend in recent decades [52]. In estuarine waters, more studies are needed to elucidate the processes and mechanisms of how exotic fishes change the original fish resources. Studies conducted at other sites in the Pearl River basin have found a direct correlation between an increase in the biomass percentage of exotic species and a reduction in fishermen’s income. Additionally, the rise in nonnative populations has been associated with a significant decrease in the functional niches of native species [39, 53]. The local fish species in the PRD can serve as essential food sources for piscivorous fish during their ontogenetic dietary shifts [54]. Despite the potential for serious economic and environmental damage resulting from the introduction and dominance of certain exotic fish species in the PRE, there is a notable lack of specific research on the associated economic and environmental impacts.

Data from the current research revealed that most nonnative fish species in this water body exhibit strong salinity adaptability. Even freshwater fish species demonstrated the ability to thrive in higher salinity waters. Field surveys indicated that salinity is a key environmental factor correlated with the spatial-temporal dynamics of fish within the PRE systems [23]. There is still a lack of research on whether indigenous fish living in estuarine waters also have strong salinity adaptability. A comparison of salinity adaptability between native and exotic fishes may help further reveal the limiting effect of salinity. However, the observed pattern of fewer nonnative fish closer to the river mouth, in addition to highlighting the significant limiting effect of salinity, may suggest that larger watershed areas and higher habitat diversity facilitate the accommodation of a greater number of nonnative fish species.

In terms of trophic levels, in the estuarine areas of large temperate rivers, the common detritivorous, piscivorous, zoobenthivorous, and zooplanktivorous fishes are generally 1-2 species, while the corresponding common fishes in the estuary areas of large subtropical rivers are usually 4-10 species [10]. Although the nutritional level of nonnative fishes upstream and downstream of the estuary does not change significantly, it is evident that the downstream waters of the estuary have a higher trophic level for nonnative fishes.

The Pearl River Basin is home to a total of 501 fish species, with the order Cypriniformes comprising 353 species, accounting for approximately 70.46% of all fish species [55]. In contrast, a fisheries resource survey along the northern coast of the South China Sea identified a total of 1,105 fish species, with up to 958 species in the PRE [56]. Among these, the order Perciformes dominates with 53.55%, while Cypriniformes represents only 2.71%. The classification structure of invasive fish species in the PRD waters shows a relatively equal proportion between Cypriniformes and Perciformes. This is evidently different from the freshwater river fish found in the Pearl River and also differs from the marine fish found in the estuary. Recent surveys conducted at eight river mouths found that during the wet season, the most abundant category was fresh migrants, with *Cirrhinus molitorella* and *Cirrhinus mrigala* being the most abundant fish species. In terms of biomass, *Oreochromis niloticus* was identified as one of the dominant specie [57]. This may suggest a higher risk of invasion into estuarine and downstream waters by more freshwater fish species, especially those with strong osmoregulatory capabilities.

The distribution of fish in the PRE is notably influenced by seasonal changes and tidal fluctuations [58]. Acoustic surveys have indicated that the overall fish density in the PRD tends to increase from upstream to the river mouth [59]. Therefore, this suggests that invasive fish species seeking to establish themselves in this region must adapt to the unique hydrodynamic patterns of estuarine waters. Field surveys focusing on the fisheries resources in the PRE reveal a lower proportion of larger-sized fish in the estuarine mouth area, but a higher density of smaller individuals [58]. For piscivorous nonnative fish species, this implies lower predation pressure and potentially abundant food resources. This could partially explain the higher trophic levels observed in invasive fish species present in estuarine waters.

One of the prominent characteristics of estuarine waters is the drastic variation in salinity, which is closely related to hydrodynamics. In terms of freshwater volume discharge, the Pearl River is ranked as the 13th largest river in the world [44]. The abundant freshwater inflow results in intense mixing of saltwater and freshwater. Hydrodynamics in the PRE (PRE) are influenced by various factors, including tidal currents, freshwater flow, wind forcing, surface heat, and coastal currents [22]. During the dry season when saltwater intrudes upstream, the surface salinity at Guadingjiao within the Modaomen Channel can increase from 2 ppt to 15 ppt within 7 hours, while the bottom salinity can decrease from 21 ppt to 1 ppt within 24 hours before rising again to 23 ppt [60]. Therefore, fish species that inhabit estuarine waters not only need to possess strong salinity tolerance but also require the ability to cope with rapid salinity changes. *Tinca tinca*, for example, has been found in tributaries but not yet in the PRD region, possibly due to its inability to adapt to the water environment in this area [61]. Investigations conducted in the lower PRE have shown that sea surface salinity is one of the crucial factors in explaining the variation in fish egg density in spawning grounds [62]. Hence, the findings of this study may hold valuable reference significance for analyzing and predicting the invasion of invasive fish species.

The results of this study reveal that even in the extreme scenario of saltwater intrusion during the rising tide, significant spatiotemporal heterogeneity persists in the salinity distribution in the PRD waters. In the eastern PRE, the pronounced temporal and spatial variation in salinity may be primarily attributed to the Pearl River plume, with the region of salinity fluctuation extending up to 40 km. As a result, brackish water covers a substantial area in the western part of the estuary, particularly in LingDing Bay [28]. This area holds the potential to become a habitat for euryhaline fish species. Long-term monitoring of this water area shows that although the range of salinity changes during flood and drought periods is similar, the average salinity is significantly different, with values of 9.95 ppt and 24.43 ppt, respectively [63]. The complex dynamics of the freshwater-saltwater interface are further compounded by the fact that the direction of the river plume is largely influenced by the monsoon and coastal currents [29]. In addition to salinity dilution, saltwater intrusion along the Modaomen Waterway is generally exacerbated by increases in tidal range and typically lasts for a maximum of 2 or 3 days [64]. Consequently, fish species invading different areas of the PRE may employ varying adaptive strategies in response to salinity changes.

For estuarine ecosystems, salinity is one of the most crucial environmental and habitat-forming factors. Investigating the impact of salinity on fish distribution in the PRE is a direct reference for studying and predicting the risk of invasive alien fish species. A comprehensive understanding of the migration behavior and habitat selection characteristics of estuarine fish is equally essential for devising appropriate survey and management strategies [65]. However, the distribution of invasive species may not be solely correlated with their adaptability to physicochemical properties of water [66]. Aquatic vegetation and covers within the habitat are equally critical for freshwater and estuarine fish because they may serve as essential sites for nesting and juvenile development [67,68]. Furthermore, the interactions among different organisms can sometimes play a significant role [10]. Nonnative fish species from other parts of the Pearl River basin would be a risk adding to their bioinvasion into the PRD [38,39]. Therefore, based on the investigation and analysis of the correlation between fish distribution and environmental factors, along with the salinity adaptability patterns of nonnative fish species studied here, and considering the biological interactions within the ecosystem, we may gain further insights into the processes and mechanisms of how the ecosystem influences the invasion of nonnative fish in the PRE. This analysis would help us understand the direct effects of invasion resistance mediated by salinity during the invasion process.

## 5. Conclusion

In summary, through the investigation, screening, and comprehensive analysis presented in this paper, a total of 25 common nonnative fish species were identified in the PRD. In terms of their use, one species is employed for biocontrol, seven for ornamental purposes, and twenty for aquaculture, with three species having both ornamental and edible value. From a phylogenetic classification perspective, these 25 nonnative fish species belong to nine different orders, with Cichliformes and Perciformes being the most predominant. Regarding water adaptability, one species is marine, while the rest are freshwater species. In terms of salinity tolerance, these nonnative fish species exhibit significant differences in their adaptability to salinity, ranging from 6.7ppt in low salinity brackish water to three times the salinity of seawater. The average salinity tolerance of the nonnative fish species is 23.74 ppt. Geographical correlation analysis indicates that the species richness of these nonnative fish decreases closer to the river mouth (p < 0.01). However, there is no significant linear change in nutritional levels from the upstream of the delta to the river mouth (p > 0.05). The investigation reveals regional and seasonal variations in salinity in the PRD. Spatial analysis demonstrates that even under extreme tidal scenarios, the salinity distribution in the PRD exhibits clear spatiotemporal heterogeneity. The area with salinity ranging from 0 to 15.0ppt covers 62.23% of the brackish water region, making it a potential habitat for the invasion of exotic fish. In conclusion, the overall analysis suggests that the majority of nonnative fish species exhibit strong salinity adaptability, emphasizing the crucial role of salinity in assessing the risk of fish invasion in estuarine environments. Further research on the patterns of salinity variation in the PRD and the salt-mediated biological interactions will contribute to screening and analyzing the invasion risk of exotic fish and elucidating the mechanisms underlying ecosystem invasibility.

## Author Contributions

Conceptualization, D.L. and D.Y.; methodology, D.L.; software, D.L.; validation, D.L., Y.Y. and Z.Z.; formal analysis, D.L.; investigation, D.L.; resources, D.L.; data curation, D.L.; writing—original draft preparation, D.L.; writing—review and editing, D.L.; visualization, D.L.; supervision, D.Y.; project administration, D.L., Y.L. and J.L.; funding acquisition, D.L., Y.L., J.L. and D.Y. All authors have read and agreed to the published version of the manuscript.

## Institutional Review Board Statement

The animal study protocol was approved by the Laboratory Animal Ethics Committee of Pearl River Fisheries Research Institute, CAFS (LAEC-PRFRI-20160323).

## Informed Consent Statement

Not applicable.

## Data Availability Statement

All data can be obtained from the authors upon request.

